# Self-Supervised Representation Learning of Protein Tertiary Structures (PtsRep) and Its Implications for Protein Engineering

**DOI:** 10.1101/2020.12.22.423916

**Authors:** Junwen Luo, Yi Cai, Jialin Wu, Hongmin Cai, Xiaofeng Yang, Zhanglin Lin

## Abstract

In recent years, deep learning has been increasingly used to decipher the relationships among protein sequence, structure, and function. Thus far these applications of deep learning have been mostly based on primary sequence information, while the vast amount of tertiary structure information remains untapped. In this study, we devised a self-supervised representation learning framework (PtsRep) to extract the fundamental features of unlabeled protein tertiary structures deposited in the PDB, a total of 35,568 structures. The learned embeddings were challenged with two commonly recognized protein engineering tasks: the prediction of protein stability and prediction of the fluorescence brightness of green fluorescent protein (GFP) variants, with training datasets of 16,431 and 26,198 proteins or variants, respectively. On both tasks, PtsRep outperformed the two benchmark methods UniRep and TAPE-BERT, which were pre-trained on two much larger sets of data of 24 and 32 million protein sequences, respectively. Protein clustering analyses demonstrated that PtsRep can capture the structural signatures of proteins. Further testing of the GFP dataset revealed two important implications for protein engineering: (1) a reduced and experimentally manageable training dataset (20%, or 5,239 variants) yielded a satisfactory prediction performance for PtsRep, achieving a recall rate of 70% for the top 26 brightest variants with 795 variants in the testing dataset retrieved; (2) counter-intuitively, when only the bright variants were used for training, the performances of PtsRep and the benchmarks not only did not worsen but they actually slightly improved. This study provides a new avenue for learning and exploring general protein structural representations for protein engineering.

## Introduction

Self-supervised learning is a powerful method for learning general representations from unlabeled samples^1-5^. In natural language processing (NLP), this takes the forms of word2vec (continuous skip-gram model, and continuous bag-of-words model)^1^, next-token prediction^2^, and masked-token prediction^3^. Recently NLP-based techniques have been applied to representation learning of protein sequences^6-10^. Two representative examples are UniRep^10^, which is based on next-token prediction and is trained on 24 million protein sequences, and a BERT^11^ model (hereinafter referred to as TAPE-BERT), which is based on masked-token prediction and is trained on 32 million protein sequences. Through transfer learning, both methods showed good performance for the prediction of protein stability landscape, and green fluorescence protein (GFP) activity landscape. For these tasks, two datasets containing about 69,000 protein sequences^12^ and 50,000 GFP variants^13^, respectively, were used. Gaining the ability to predict protein engineering outcomes is a major goal for biotechnology, which would significantly expand the uses of proteins and enzymes for pharmaceutical, industrial, and agricultural purposes. However, predictions based on traditional rational design approaches have not yet reached the level of accuracy required for routine practice^12,14-16^, and the directed protein evolution approach often demands iterative processes and high throughput screening assays^17,18^. Artificial intelligence approaches such as machine learning^16^, UniRep^10^, and TAPE-BERT^11^ have great potentials to accelerate the rational design of proteins.

The methods for protein representation learning developed thus far have mostly relied on protein sequence information, whereas the vast amount of protein tertiary structure information available in the protein data bank (PDB)^19^ remains unused, and the recent release of the breakthrough algorithm AlphaFold 2^20^ will likely make protein tertiary structures more readily available. How to utilize protein structural information in deep learning remains a fundamental open question whose answers will strongly benefit protein design and engineering.

In this work, we present a self-supervised learning model designed to extract embedded representations of protein tertiary structures (PtsRep) (Fig. 1), using KNR (*K* nearest residues) as the input format for the protein structures (Fig. 1A). Considering that in proteins the properties of a residue are affected by the surrounding residues^21^, we adopted the bidirectional language model^22^, an advanced next-token algorithm, to predict multiple neighboring residues on both forward and backward directions of a given protein. In order to increase the stringency of the prediction, the residues immediately adjacent to any given residue were omitted from the prediction. The learned representations summarized the protein tertiary structures into fixed-length vectors of 768 dimensions. These vectors were then used for two tasks: (1) prediction of protein stability, and (2) prediction of GFP fluorescence brightness (Fig. 1C). The prediction performances of PtsRep were compared to those of the benchmarks UniRep and TAPE-BERT. In addition, the advantages of PtsRep over the benchmarks was evaluated in terms of “testing budget” required to identify a target variant, which has critical implications in the application of these machine learning methods to answer practical protein engineering questions.

**Figure 1.**
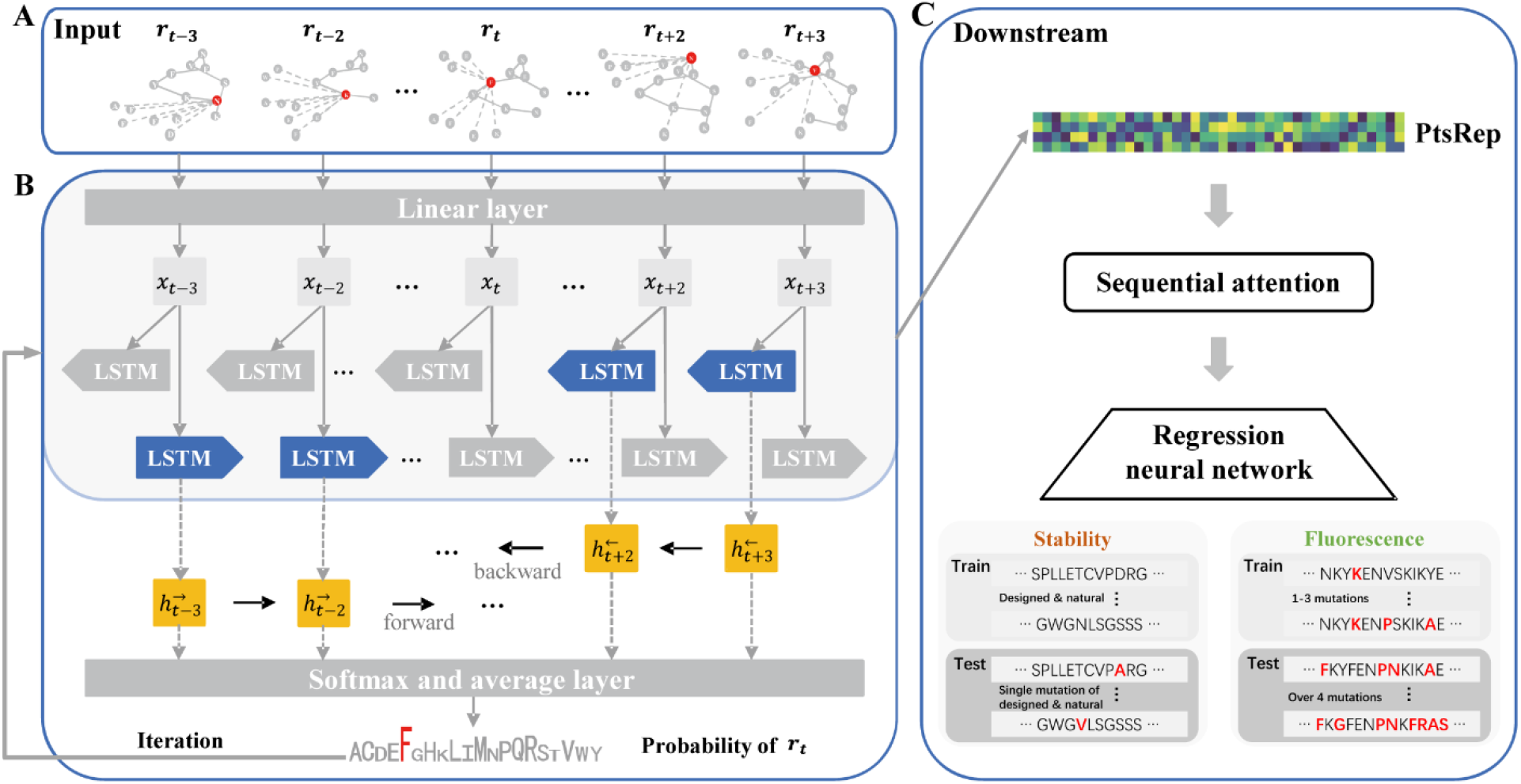
Workflow used by PtsRep for learning and applying protein tertiary structure representation. (A) In the input module, protein structures from PDB were encoded with the KNR algorithm, with each amino acid represented by the nearest 15 amino acids and their features. (B) In the learning (pre-training) module, PtsRep performed contextual noncontiguous residue prediction using a cross-entropy loss function, and internally represent proteins. (C) A regression neural network with attention mechanism was used to transfer the embedded representations to downstream prediction tasks.

## Methods

### 1. Representation learning network

The learning network architecture is shown in Fig. 1 and Supplementary Fig 1. The bidirectional language model ^2,22^ was used to predict the two contiguous residues beside any given residues in both directions, with a ***d*** number of adjacent residues omitted. Specifically, we defined *R(****t, d***, the sum of log likelihood in both the forward and backward directions, as follows:

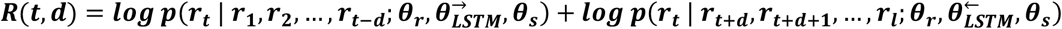

Where, ***r*** refers to a residue (token) in an input protein tertiary structure, ***l*** as the length of a protein, ***t*** as the position of the residue in the protein sequence. *θ*_***x***_ as the *K* nearest residue representations of a protein, *θ*_***LSTM***_ as the LSTM layer, *θ*_***s***_ as the Softmax layer, and ***p*** as the probability^23^ of the predicted residue corresponding to the actual residue.

We then calculated *R(****t, d***) for every ***t*** ∈ {**1, 2**, …, ***l***} and ***d*** ∈ {***d, d*** + **1**}, where the instances ***t*** − ***d*** < 0 and ***t*** + ***d*** > ***l*** were not calculated. The final loss was the average of the calculated ***Loss*** for all positions of ***t***.

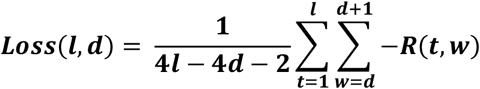

Considering the large amount of protein structural data used in this study, we adopted a double-layer bi-LSTM, and chose 768 as the dimension of embeddings, as in TAPE-BERT^11^. Convergence was defined as no improvement in the validation loss for 5 epochs. The best model was trained for 1.8 million weight updates corresponding to ∼50 epochs. Because the length of each protein sequence was different, we used a batch size of 1. All models were trained with the Adam optimizer^24^. Where indicated, Swish activation^25^ and layer normalization^26^ were applied, and to reduce the risk of overfitting, a dropout rate of 10% was applied to the output layer (supplementary Fig. 1).

### 2. Benchmark representations

We used UniRep, which is an mLSTM model trained on about 24 million protein sequences, as a benchmark representation^10^. We downloaded the trained weights and obtained the representations using the code in https://github.com/churchlab/UniRep. We chose the 1900-dimensional UniRep representation, which was reported to perform best among the different dimensions^10^. We also used TAPE-BERT, which is trained on about 32 million protein sequences, as an additional benchmark representation^3^. We downloaded the trained weights and obtained the representations using the code in https://github.com/songlab-cal/tape. Furthermore, we used one-hot encoding and KNR as baseline representations.

### 3. Downstream task evaluation

To better compare the performances with the benchmarks, we used the same downstream architecture with attention mechanism and multilayer perceptron, as described in TAPE-BERT^11^. The embeddings of protein tertiary structures extracted from the pretrained network were used without end-to-end optimization. For these tasks, we reported the Pearson’s ***γ*** (product-moment correlation coefficient), Spearman’s *ρ* (rank correlation coefficient), accuracy (ACC) and mean-square error (MSE) from different representations. Five-fold random cross validation was used, and the 10 models with the highest Spearman’s *ρ* on the validation dataset of each fold were selected as the test mode. Convergence was defined as an improvement lower than 0.002 for the Spearman’s *ρ* after 20 epochs during the validation.

### 4. Datasets

#### 1) Self-supervised learning dataset

We used ProteinNet^27^ as the training and validation sets for the self-supervised learning model. We first collected the 90% thinning version of ProteinNet12^27^, which included 49,600 protein chains with tertiary structures. Then we excluded the protein chains (1) with missing Cα coordinates, or (2) with sequences longer than 700 aa or shorter than 30 aa. The resulting dataset had 35,568 protein chains. We used 95% of the data as the training dataset, and 5% as the validation dataset (Supplementary Table 1).

#### 2) Stability landscape prediction dataset

We used the dataset from Rocklin *et al*^12^. This set includes the sequences and the chymotrypsin stability scores of 69,034 proteins (17,773 *de novo* designed proteins, 10,674 variants of these designed proteins, 1,193 natural proteins, 2,423 variants of these natural proteins, 24,900 scrambled versions of the proteins, and 12,071 control inactivated sequences where a buried aspartate residue was inserted). Among these proteins or variants, we used a total of 16,431 proteins or variants with structural information (16,159 *de novo* designed mini proteins and 272 natural proteins) for training and validation, and 12,851 point variants of 14 de novo designed proteins and 3 natural proteins with structural information for testing^12^. For training and validation, we used an 80%/20% splitting strategy (Supplementary Table 1).

#### 3) Fluorescence landscape prediction

We used the dataset from Sarkisyan *et al*^13^, which contains more than 50,000 variants of green fluorescent protein from *Aequorea victoria*. The number of amino acid substitutions in the variants ranged from 1 to 15^14^. Following a previous study, we used the variants with 1∼3 substitutions (a total of 26,198 variants) for training and validation, and the variants with 4 or more substitutions (a total of 25,517 variants) for testing^20^. For training and validation, we used an 80%/20% splitting strategy (Supplementary Table 1).

## Results

### 1. Network architecture for protein tertiary structure representation learning (PtsRep)

PtsRep comprises of three modules as shown in Fig. 1. In the input module (Fig. 1A), to feed the protein tertiary structural information, we adopted an algorithm^28^ that enables each residue to be represented by a series of properties (i.e., bulkiness^29^, hydrophobicity^30^, flexibility^31^, relative spatial distance, relative sequential residue distance, and spatial position based on spherical coordinate system) of its *K* nearest residues (KNR) in the Euclidean space^28^. The parameter *K* was set at 15 after initial optimization based on the performance criteria for the downstream tasks (supplementary Table 2). A dataset of 35,568 protein chains was selected from PDB, 95% of which was used for training and 5% for validation. In the training module (Fig. 1B), a bidirectional language model^2^ was adopted to predict the four nearest noncontiguous residues on both the forward and backward directions of a given amino acid ***t***, with the immediate adjacent residue omitted (*i*.*e*., positions ***t*** - **3, *t*** - **2, *t*** + **2, *t*** + **3**). The omission of these two residues was again chosen after optimization based on the performance criteria for the downstream tasks (see supplementary Table 3, and Methods). The network was iterated to maximize the prediction accuracy. This yielded a top model, which summarized the protein tertiary structures into ***l*** × 768-dimensional vectors for any given protein sequence of length ***l***. Finally, this trained or pre-trained (with respect to the downstream tasks) PtsRep model was applied to two protein landscape prediction tasks, protein stability and green fluorescence protein brightness, through a downstream network (se Methods)^11^, as shown in Fig. 1C.

### 2. PtsRep significantly outperformed benchmarks on protein stability prediction

Protein stability is a key objective for protein engineering. In this study, we used the dataset from Rocklin *et al*^12^, which consists of the following four protein topologies: *ααα, αββα, βαββ, ββαββ*, where *α* denotes a α-helix and *β* denotes a β-sheet. It should be noted that the training/validation dataset for the benchmarks UniRep and TAPE-BERT contains 56,180 proteins, among which only 16,431 have structural information suitable for PtsRep processing (Supplementary Table 1). Thus, the PtsRep model was trained on only 29.3% of the data used for UniRep and TAPE-BERT. However, all the proteins in the testing dataset for the benchmarks had solved structures, thus the PtsRep model was tested on the same testing dataset used in previous studies for UniRep and TAPE-BERT.

As shown in Table 1, we compared Spearman’s ***ρ***, ACC and MSE. We found that PtsRep significantly outperformed the best benchmark TAPE-BERT (Spearman’s *ρ* = 0.79 *versus* 0.73), and much more so the baseline KNR (Spearman’s *ρ* = 0.79 *versus* 0.37). PtsRep also generally showed better ACC and MSE, compared with both the benchmarks. Interestingly, the benchmark UniRep performed differently for different protein topologies (e.g., *αββα, ααα*), whereas PtsRep performed more evenly in this regard, and resulted in a low standard deviation, similar to that of TAPE-BERT (0.11 *versus* 0.08), but significantly better than that of UniRep (0.11 *versus* 0.25) (Supplementary Table 4).

**Table 1.**
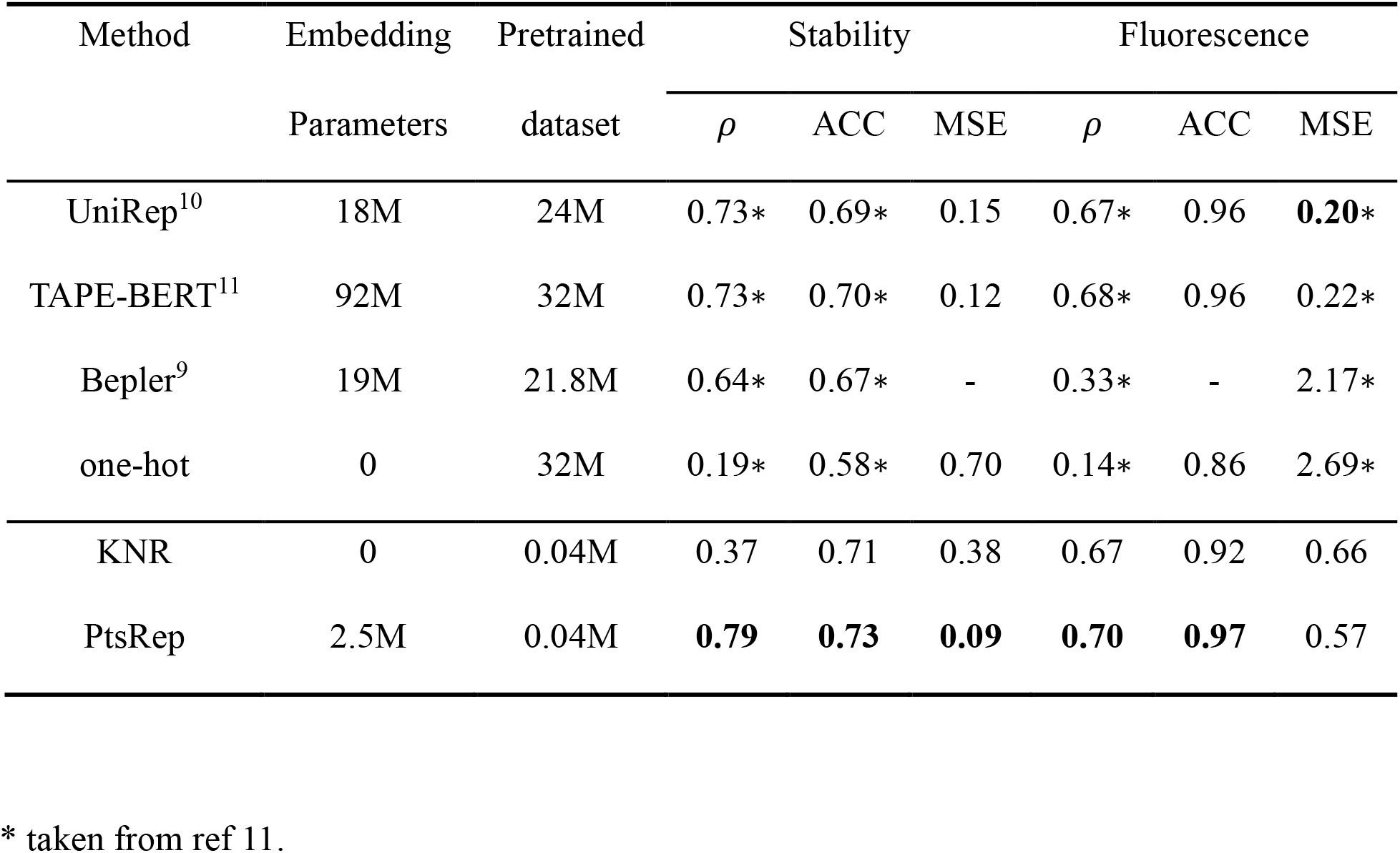
Embedding performances for downstream tasks of PtsRep, the outlined benchmarks and the baselines.

### 3. PtsRep improved the prediction of the fluorescence brightness of GFP variants

Since the protein stability landscape dataset consists of mainly artificially designed small proteins and only an insufficient number of natural proteins, in further endeavors we focused on the GFP variants dataset of Sarkisyan *et al*^13^. As described in a previous study^11^, we allocated the variants with 1-3 substitutions (for a total of 26,198) as the dataset for training and validation, and the variants with four or more substitutions (for a total of 25,517) as the dataset for testing. As shown in Table 1, PtsRep outperformed the best benchmark (*i*.*e*., TAPE-BERT) in this task (Spearman’s *ρ* = 0.70 *versus* 0.68).

Since these statistical metrics are often difficult to correlate with the actual practice of protein engineering, we explored an alternative and more direct measure, the “testing budgets” for target variants^10,11^. We selected the top 0.1% of the GFP variants in the testing dataset in terms of brightness, for a total of 26 variants (with 4 to 15 mutations), and tested the ability of each method to prioritize this set of brightest variants under different testing budgets. As shown in Fig. 2A, PtsRep performed better than the benchmarks UniRep and TAPE-BERT, and significantly better than the baselines. For example, PtsRep achieved a recall rate of 70% for this top 0.1% brightest variants with about 380 variants retrieved, compared with 559 variants for TAPE-BERT, 737 variants for UniRep, and 1,609 variants for the baseline KNR, or 1.5-fold, 1.9-fold, and 4.2-fold the experimental burden required for PtsRep, respectively.

**Figure 2.**
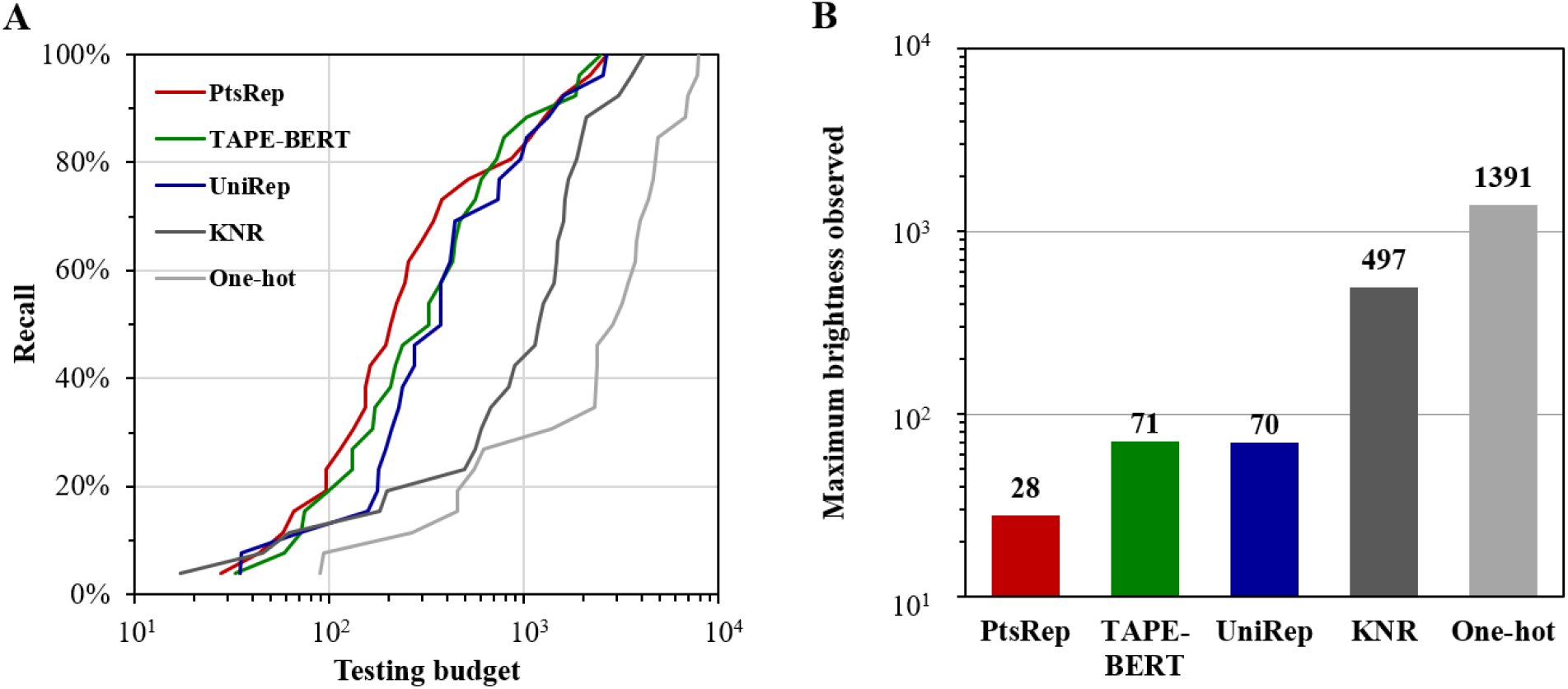
(A) Recall rates of the top 0.1% brightest GFP variants retrieved *versus* the required testing budget for each representation method. (B) The number of trials required to identify the brightest GFP variant based on the ranking obtained with each representation method.

Subsequently, we tested how the brightest GFP variant fared in the ranking obtained with each method^10,11^. As shown in Fig. 2B, we found that the brightest protein ranked 28^th^ when using PtsRep, whereas it ranked 71^st^ when using TAPE-BERT, respectively, and 497^th^ for the baseline KNR, corresponding to 2.5-fold and 17.8-fold the experimental cost for PtsRep, respectively.

### 4. Refinement of PtsRep for the prediction of the fluorescence brightness of GFP variants

We then asked two additional questions: (1) what if only a fraction of the training dataset was used? and (2) what if only the bright mutants were used in the training dataset?

As shown in Supplementary Fig. 2, the use of a subset representing 20% of the whole training and validation dataset (randomly selected, for a total of 5,239 variants) resulted in a fairly satisfactory performance. In particular, PtsRep achieved a recall rate of 70% for the top 26 brightest variants with about 795 variants retrieved, *i*.*e*., only 2-fold the experimental burden required when using the whole training dataset. The brightest GFP variant ranked 87^th^, *i*.*e*., 3-fold the experimental burden required when using the whole training dataset. For comparison, TAPE-BERT needed to retrieve 1,347 variants to reach the same recall rate, and the brightest GFP variant ranked 119^th^.

Very surprisingly, however, when the dark variants were removed from the training dataset (resulting in a total of 21,316 variants), the prediction performances of all methods did not suffer, but rather slightly improved. For example, to achieve a recall rate of 70% for the top 26 brightest variants, PtsRep and TAPE-BERT now needed to retrieve a reduced number of 314 and 432 variants, respectively. Furthermore, the brightest GFP variant ranked 7^th^ and 6^th^, respectively (Supplementary Fig. 3). When 20% of these bright variants (randomly selected, for a total of 4,263 variants) was used, to obtain the same recall rate, PtsRep and TAPE-BERT needed to retrieve 761 and 926 variants, respectively, and the brightest GFP variant ranked 84^th^ and 88^th^, respectively.

**Figure 3.**
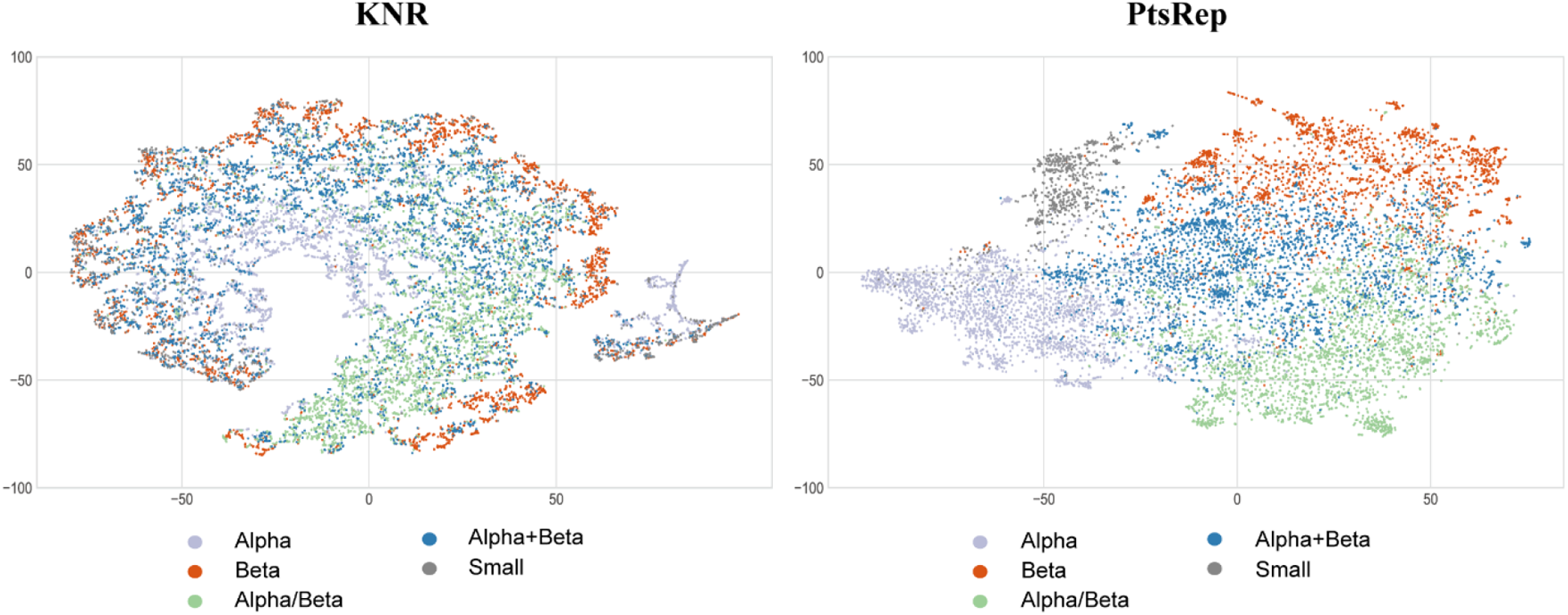
t-SNE representations obtained with KNR (left panel) and PtsRep (right panel) for 15,444 proteins classified by the Structural Classification of Proteins (SCOP)^34^. The t-SNE projections from the embedded space onto a low dimensional representation are shown. The sequences from SCOP are colored according to their ground-truth structural class (alpha, beta, alpha/beta, alpha+beta, and small proteins).

Lastly, ensemble methods with weighted contributions of multiple models are known to generally improve the performance of machine learning models^32^. In our case, a combination of the three different representation methods PtsRep, TAPE-BERT, and UniRep improved the prediction performance as expected. As shown in supplementary Table 5, for example, to reach a recall rate of 70% for the top 0.1% brightest variants, the PtsRep-TAPE-BERT-UniRep model needed to retrieve 329 variants compared with 380 variants for the PtsRep model alone, and the brightest GFP variant ranked 21^st^ with the combined model, compared with 28^th^ for the PtsRep model alone.

### 5. PtsRep performed well for protein fold classification

To examine what PtsRep learned from the PDB^19^ dataset, we used t-distributed stochastic neighbor embedding (t-SNE)^33^ to test its unsupervised clustering ability on a dataset of 15,444 semantically related unlabeled proteins, which have been classified in the Structural Classification of Proteins (SCOP)^34^ database. We found that PtsRep well separated the five different types of protein structures (Fig. 3, right panel). Using the Davies-Bouldin Index (DBI)^35^, a general index for evaluating clustering algorithms, we found that the DBI for PtsRep was as low as 1.3. In contrast, for the baseline representation KNR, which utilized the same amount of structural information but was not trained with PtsRep, the DBI was 11.6 (Fig. 3, left panel). Therefore, the trained PtsRep model improved the clustering by 8.9-fold in this SCOP classification task.

We also used t-SNE to test the ability of PtsRep to cluster the GFP variants. As shown in supplementary Fig. 4, the variants with bright fluorescence were more clustered when using PtsRep resulting in a DBI index of 2.5, compared with the more scattered distribution obtained when using KNR, which scored a DBI of 4.6.

## Discussion

In this work, we presented PtsRep, a self-supervised learning method that was designed to learn general protein structure representations from unlabeled protein tertiary structures deposited in the PDB archive. The learned embeddings were applied to the predictions of protein stability and GFP fluorescence using two publicly available datasets. PtsRep showed an outstanding performance on both tasks compared with the protein sequence-based benchmark methods UniRep and TAPE-BERT. Compared to the best benchmark TAPE-BERT, which is trained on 32 million protein sequences (with 96 million parameters), our PtsRep model was trained on only 35,568 protein structures (with 2.5 million parameters), or 0.11% of the data used in TAPE-BERT in terms of protein entries.

While what deep learning learns from data is often considered a black box, we attempted a partial dissection of what PtsRep has learned from the PDB dataset. Given the fact that PtsRep was able to perform much better than KNR, in terms of protein fold classification, as shown in Fig. 3, we suggest that PtsRep has at least understood which protein structures are stable, since unstable and thus inactive protein structures are not likely present in the PDB. We also know from the practice of directed evolution of enzymes that around 30% of randomly mutated protein variants are inactive^36^. This explains, in part, the performance improvement that we saw for PtsRep over UniRep and TAPE-BERT as shown in Fig. 2.

This same reason could be a major contributor to the convergence of the performances of the benchmarks towards that of PtsRep when only bright GFP variants were used for training (Fig. 3), as if in this manner the benchmarks indirectly gained the ability to distinguish stable bright GFP variants as well as PtsRep. Nonetheless, it is worth noting that even in such instance PtsRep was still superior to Unirep and TAPE-BERT, since it was trained on only a very small fraction of the data used for the benchmarks.

However, how PtsRep is able to cluster GFP variants better then KNR remains to be understood (supplementary Fig. 2). Since a similar observation was made for TAPE-BERT^11^, which is trained on protein sequences alone, it is likely that the better clustering of GFP seen for PtsRep was not only structure-driven, but also partially sequence- and algorithm-driven. Along this line, it is interesting to note that the performance of PtsRep was still much better than the baseline model KNR, and was in fact close to those of UniRep and TAPE-BERT even when the parameter *K* was set at 1 (supplementary Table 2). This can likely be attributed to two aspects of the learning algorithm: 1) the fact that the algorithm predicts not just one “next token,” but two continuous residues in both directions, 2) the fact that the algorithm omits the adjacent residues that might be easy to predict given the KNR format, both of which might force the model to better internally represent proteins (supplementary Table 3). Predicting more than two continuous residues, however, failed to further improve the performance of PtsRep for downstream tasks (data not shown).

As far as protein engineering is concerned, we have learned two very important lessons from the deep learning of GFP fluorescence landscape: 1) only a few thousand variants are sufficient to train a predictive model, which is experimentally manageable; and 2) the active (*i*.*e*., bright) variants provide a better guidance. If the latter holds true for other proteins, this could result in a tremendous saving in terms of experimental cost, as it would not be necessary to characterize inactive variants. Lastly, it is also possible to combine PtsRep (alone or together with TAPE-BERT and UniRep) with traditional methods such as simulated annealing^37^ and Bayesian optimization^38^, and extend the use of these deep learning strategies for disease diagnosis^10,39,40^, and drug design^41,42^.

## Acknowledgments

This work was supported by the National Key R&D Program of China (2018YFA0901000), and the Guangzhou Science and Technology Program key projects (201904020016). We thank Xing Zhang for technical assistance.

