## Supplemental Figures 1, 2, 3 and 4; Supplemental Tables 1, 2, 3, 4 and 5. for "Self-Supervised Representation Learning of Protein Tertiary Structures (PtsRep) and Its Implications for Protein Engineering"

### Supplementary Materials

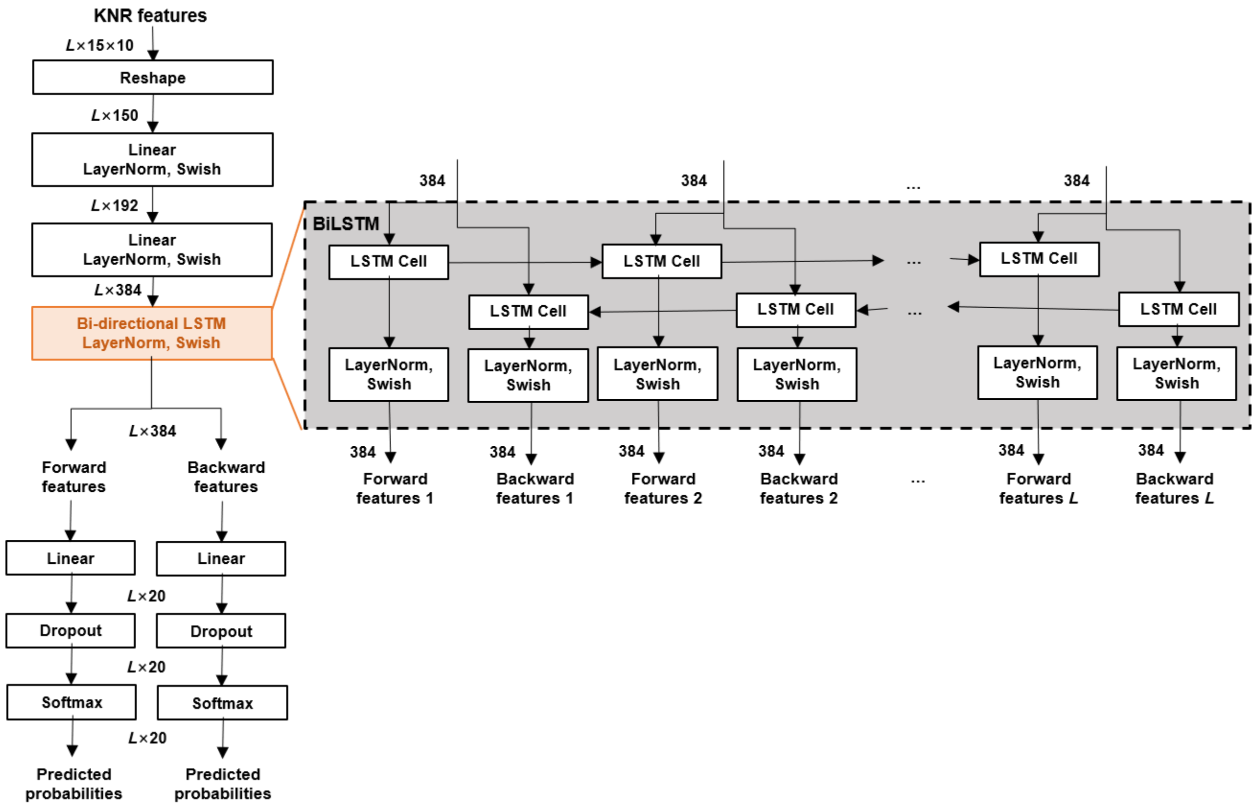


**Supplementary Figure 1.** Network architecture of the learning (pre-training) module of PtsRep. Bi-LSTM was used to extract the forward and backward contextual information. The Feedforward Neural Network was used to change dimension and predict the types of amino acids at different positions. LayerNorm, layer normalization[1]; Swish, swish activation[2].

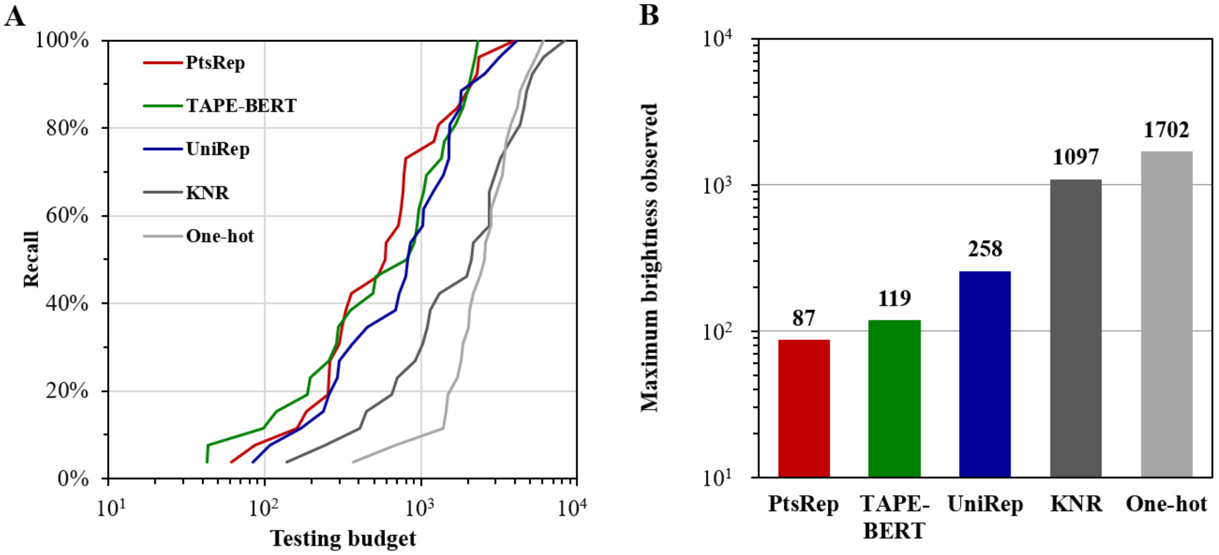


**Supplementary Figure 2. Effect of a reduced training set (20% of the original dataset) of variants on the prediction of GFP fluorescence brightness.** (A) Recall rates of the top 26 brightest GFP variants retrieved *versus* the required testing budget for each representation method. (B) The number of trials required to identify the brightest GFP variant based on the ranking obtained with each representation method.

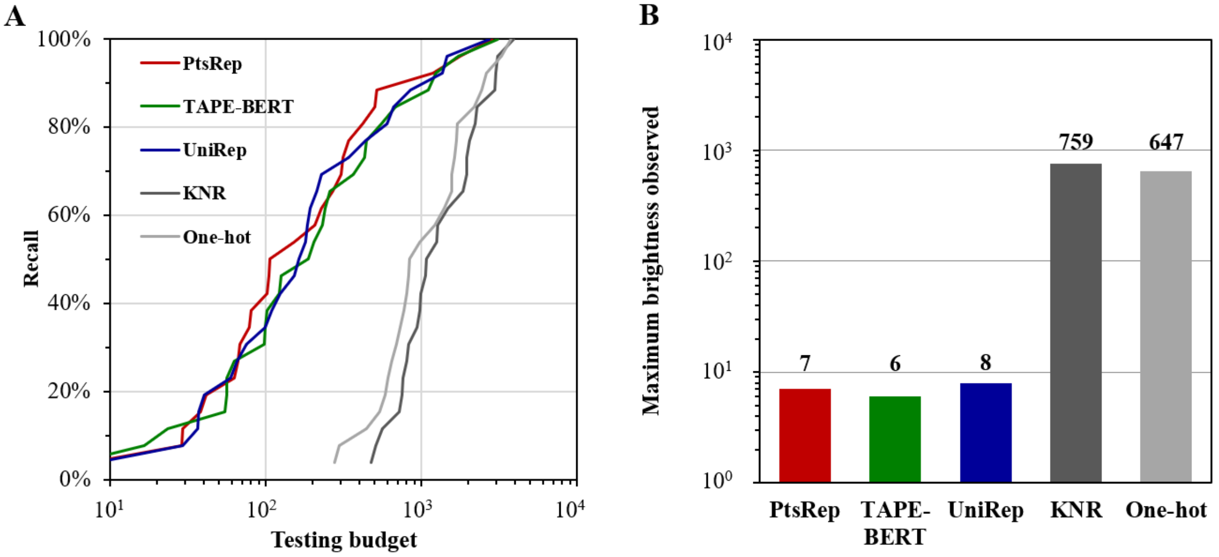


**Supplementary Figure 3. Effect of using a training dataset including only bright variants on the prediction of GFP fluorescence brightness.** (A) Recall rates of the top 26 brightest GFP variants retrieved *versus* the required testing budget for each representation method. (B) The number of trials required to identify the brightest GFP variant based on the ranking obtained with each representation method.

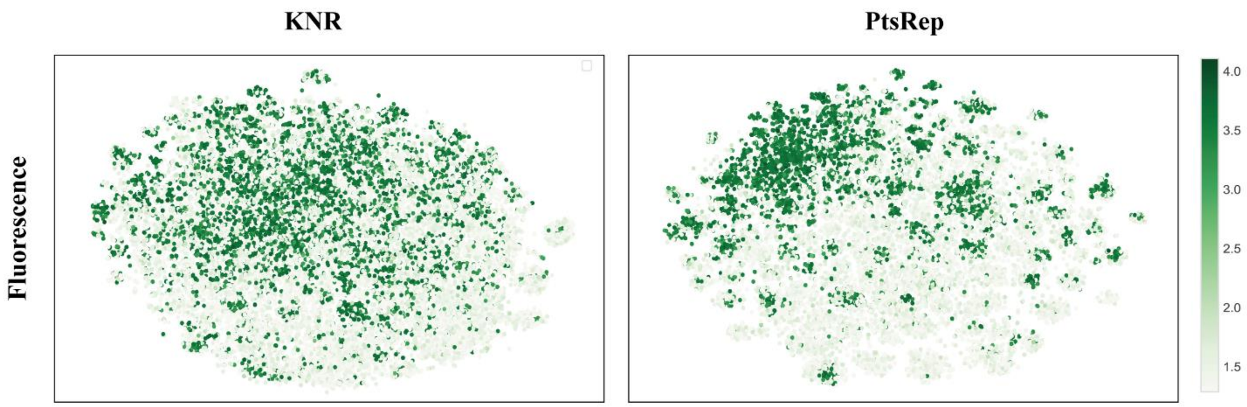


**Supplementary Figure 4.** t-SNE visualization obtained with KNR (left panel) and PtsRep (right panel) for GFP variants with 4 to 15 substitutions.

**Supplementary Table 1.** Tertiary structure learning (pretraining) and downstream tasks dataset sizes.

| Dataset | Training set size | Validation set size | subtotal | Test set size |
|---|---|---|---|---|
| Pretraining[3] | 33,790 | 1,778 | 35,568 | - |
| Protein stability[4] | 44,901 | 11,225 | 56,126 | 12,851 |
| with structure | 13,145 | 3,286 | 16,431 | 12,851 |
| GFP fluorescence brightness[5] | 20,958 | 5,240 | 26,198 | 25,517 |
| bright variants | 17,052 | 4,264 | 21,316 | 8,249 |
| dark variants | 3,906 | 976 | 4,882 | 17,268 |

**Supplementary Table 2.** The performances for protein encoding with different K nearest residues in the input module.

| K Nearest | Stability | | | | Fluorescence | | | |
|---|---|---|---|---|---|---|---|---|
| | $\gamma$ | $\rho$ | ACC | MSE | $\gamma$ | $\rho$ | ACC | MSE |
| 1 | 0.70 | 0.67 | 0.72 | 0.16 | 0.79 | 0.69 | 0.96 | 0.56 |
| 5 | 0.75 | 0.75 | **0.73** | 0.12 | 0.81 | 0.69 | 0.96 | 0.56 |
| 10 | 0.77 | 0.77 | 0.71 | 0.14 | **0.82** | 0.69 | 0.96 | **0.54** |
| 15 | **0.78** | **0.79** | **0.73** | **0.09** | **0.82** | **0.70** | **0.97** | 0.57 |
| 20 | 0.74 | 0.74 | 0.73 | 0.13 | 0.80 | 0.70 | 0.96 | 0.62 |
| 25 | 0.72 | 0.73 | 0.73 | 0.13 | 0.80 | 0.69 | 0.96 | 0.56 |

**Supplementary Table 3.** Embedding performances for different predicted positions.

| Predicted position | Stability | | | | Fluorescence | | | |
|---|---|---|---|---|---|---|---|---|
| | $\gamma$ | $\rho$ | ACC | MSE | $\gamma$ | $\rho$ | ACC | MSE |
| -2, -1, 1, 2 | 0.68 | 0.65 | **0.74** | 0.13 | 0.78 | 0.69 | 0.96 | 0.70 |
| -3, -2, 2, 3 | **0.78** | **0.79** | 0.73 | **0.09** | **0.82** | **0.70** | **0.97** | 0.57 |
| -4, -3, 3, 4 | 0.73 | 0.72 | 0.73 | 0.12 | 0.82 | 0.69 | 0.96 | **0.50** |
| -1, 1 | 0.56 | 0.52 | 0.72 | 0.18 | 0.76 | 0.60 | 0.96 | 0.71 |
| -2, 2 | 0.67 | 0.68 | 0.73 | 0.13 | 0.78 | 0.70 | 0.96 | 0.63 |

**Supplementary Table 4.** PtsRep performances across 4 topologies in the stability prediction task.

| Method | $\alpha\alpha\alpha$ | | $\alpha\beta\beta\alpha$ | | $\beta\alpha\beta\beta$ | | $\beta\beta\alpha\beta\beta$ | | Test dataset | | |
|---|---|---|---|---|---|---|---|---|---|---|---|
| | $\rho$ | ACC | $\rho$ | ACC | $\rho$ | ACC | $\rho$ | ACC | $\rho$ | ACC | Std |
| UniRep* | 0.72 | 0.66 | 0.11 | 0.76 | **0.68** | 0.66 | 0.65 | 0.67 | 0.73 | 0.69 | 0.25 |
| TAPE-BERT* | 0.66 | **0.68** | 0.48 | 0.73 | 0.65 | **0.71** | 0.65 | 0.67 | 0.73 | 0.70 | **0.08** |
| Bepler* | 0.33 | 0.66 | 0.24 | **0.79** | 0.54 | 0.70 | 0.58 | 0.53 | 0.64 | 0.67 | 0.14 |
| One-hot* | 0.58 | 0.59 | 0.04 | 0.58 | -0.05 | 0.58 | 0.54 | 0.58 | 0.19 | 0.58 | 0.29 |
| KNR | 0.54 | 0.63 | 0.39 | 0.77 | -0.05 | 0.70 | 0.32 | 0.69 | 0.36 | 0.71 | 0.22 |
| PtsRep | **0.73** | 0.64 | 0.44 | **0.80** | 0.49 | 0.64 | 0.64 | **0.75** | **0.79** | **0.73** | 0.11 |

* taken from ref 6.

**Supplementary Table 5.** Recall Performances for PtsRep, PtsRep-TAPE-BERT-UniRep, and TAPE-BERT-UniRep (for the top 26 variants, and the brightest variant, with recall rates of 70%)

| Method | the top 26 variants | the brightest variant ranked |
|---|---|---|
| One-hot | 4362 | 1391 |
| KNR | 1609 | 497 |
| UniRep | 737 | 70 |
| TAPE-BERT | 559 | 71 |
| PtsRep | 380 | 28 |
| TAPE-BERT-UniRep (50%/50%) | 378 | 39 |
| PtsRep-TAPE-BERT-UniRep (50%/25%/25%) | **329** | **21** |
